# Genomic evidence for a hybrid origin of the yeast opportunistic pathogen *Candida albicans*

**DOI:** 10.1101/813436

**Authors:** Verónica Mixão, Toni Gabaldón

## Abstract

Opportunistic yeast pathogens are of increasing medical concern. *Candida albicans*, the species with the highest incidence, is a natural commensal of humans that can adopt a pathogenic behaviour. This species is highly heterozygous, is an obligate diploid, and cannot undergo meiosis, adopting instead a parasexual cycle. The origin of these traits is unknown and we hypothesize they could result from ancestral hybridization. We tested this idea by analyzing available genomes of *C. albicans* isolates and comparing them to those of hybrid and non-hybrid strains of other *Candida* species. Our results show compelling evidence that *C. albicans* is an evolved hybrid, with levels and patterns of ancestral heterozygosity that cannot be fully explained under the paradigm of vertical evolution. Although the level of inferred divergence between the putative parental lineages (2.8%) is not clearly beyond current species boundaries in Saccharomycotina, we show here that all analyzed *C. albicans* strains derive from a single hybrid ancestor, which diverged by extensive loss of heterozygosis. This finding has important implications for our understanding of *C. albicans* evolution, including the loss of the sexual cycle, the origin of the association with humans, and the evolution of virulence traits.

## Introduction

*Candida* species are the most common causative agents of hospital-acquired fungal infections (Pfaller and Diekema 2007; Lass-Flörl 2009; Brown et al. 2012; Gabaldón and Carreté 2016; OPATHY Consortium and Gabaldón 2019), accounting for 72.8 million opportunistic infections per year, with an overall mortality rate of 33.9% (Jordà-Marcos et al. 2007; Pfaller and Diekema 2007). *Candida albicans* is a commensal organism that can form part of the microbiota of healthy individuals (Cauchie, Desmet, and Lagrou 2017). Under certain circumstances, such as a weakening of the host immune system, *C. albicans* can shift from a commensal to a pathogenic behaviour (Pfaller and Diekema 2007). This species is the causative agent in more than 50% of the candidaemia cases worldwide (Pfaller and Diekema 2007). Although *C. albicans* cannot undergo a normal sexual cycle involving meiosis, it is known to be able to follow a so-called parasexual cycle (Forche et al. 2008; Berman and Hadany 2012). This consists of the mating of two diploid cells, forming a tetraploid cell that subsequently returns to a diploid state by concerted chromosomal loss. Restoration of the diploid state is not always properly achieved, leading to aneuploidies (Berman and Hadany 2012; Bennett 2015). Thus, this system constitutes a source of genetic variability, which has been proposed to be advantageous under stress conditions (Berman and Hadany 2012).

Recently, the genomic diversity of *C. albicans* strains belonging to different MLST-based clades and isolated from globally distributed locations was investigated (Hirakawa et al. 2015; Ene et al. 2018; Ropars et al. 2018; Wang, Bennett, and Anderson 2018; Bensasson et al. 2019). These studies have shown that *C. albicans* genome shows signs of recombination, with genomic material exchanged between different strains (Ropars et al. 2018; Wang, Bennett, and Anderson 2018). Furthermore, they have reported that the fraction of the genome covered by heterozygous regions can vary between 48% to 89%, depending on the strain, and that these heterozygous tracts are separated by regions of loss of heterozygosity (LOH) (Ene et al. 2018; Ropars et al. 2018; Wang, Bennett, and Anderson 2018; Bensasson et al. 2019). Moreover, it has been shown that the accumulation of mutations and the exchange of genomic material between strains are the main forces shaping *C. albicans* genome (Wang, Bennett, and Anderson 2018).

However, an intriguing and still unaddressed question is: what is the initial source of the high heterozygosity levels present in *C. albicans* genome? Can the accumulation of mutations over long periods of time, and the presence of inter-strain recombination explain the levels of heterozygosity observed in highly heterozygous regions of *C. albicans* strains? We noted that the genomic patterns observed in *C. albicans* are reminiscent of recently analyzed yeast hybrid species, such as *Candida inconspicua, Candida orthopsilosis* and *Candida metapsilosis* (Pryszcz et al. 2014; Pryszcz et al. 2015; Schröder et al. 2016; Ropars et al. 2018; Mixão et al. 2019). Hybrids are chimeric organisms originated from the hybridization between two diverged lineages (whether from the same or distinct species), and consequently have genomes with high levels of heterozygosity. Diploid hybrids comprise pairs of homeologous chromosomes, which divergence at the sequence level is initially similar to the one of the two parentals at the time of hybridization (Mixão and Gabaldón 2018). These genomes can subsequently evolve through recombination-mediated conversion of homeologous sequences, resulting in LOH (Mixão and Gabaldón 2018). Hybridization may originate organisms with unique phenotypic features, which may contribute to the successful adaptation to new niches (Gladieux et al. 2014). Finally, hybridization has been recently hypothesized to be at the root of the origin of the virulent potential in some emerging yeast pathogens, such as *C. metapsilosis* or *C. inconspicua* (Pryszcz et al. 2015; Mixão and Gabaldón 2018; Mixão et al. 2019). Hence, a possible scenario for the observed genomic patterns in *C. albicans* is that the divergence observed in highly heterozygous regions is not exclusively the consequence of continuous accumulation of mutations within the lineage, but also, to a large degree, a footprint of an ancient hybridization event between two diverged lineages. We here put these alternative hypotheses at test by comparing *C. albicans* genomic patterns with the ones observed in *C. inconspicua, C. orthopsilosis* and *C. metapsilosis* hybrid strains, as well as, non-hybrid strains from these and other species.

## Results

### *K*-mer profiles of *C. albicans* sequencing libraries reveal a heterogeneous content similar to that of hybrid genomes

To assess heterozygosity levels in *C. albicans* genomes, we analyzed 27-mer frequencies of raw sequencing data of Illumina paired-end libraries from different *C. albicans* strains (see Materials and Methods). All analyzed sequencing libraries produced similar 27-mer profiles, showing two peaks of depth of coverage, one with half coverage of the other, which corresponded to heterozygous and homozygous regions, respectively (Figure 1A and Figure S1). Of note, for all strains, including the reference, approximately half of the 27-mers of the first peak were not represented in the reference assembly (Figure 1A and Figure S1), and therefore could correspond to heterozygous regions where only one of the haplotypes was represented in the reference assembly. This bimodal pattern was also produced by sequencing libraries from hybrid strains of *C. orthopsilosis* and *C. metapsilosis* mapped to their respective reference assemblies (Figure 1A and Figure S1) and was previously reported for *C. inconspicua* hybrids (Mixão et al. 2019). However, this pattern was not observable in sequencing libraries from non-hybrid strains of *Candida dubliniensis, Candida tropicalis, C. orthopsilosis* and *Candida parapsilosis* (Figure 1A and Figure S1). As shown in Figure S1, hybrid strains with lower levels of heterozygosity (i.e. strains from *C. orthopsilosis* clade 1) had a higher homozygous peak as compared to hybrid strains with higher levels of heterozygosity (i.e. *C. orthopsilosis* clade 4). Altogether, these results show that the relative frequency of the two peaks is representative of the level of heterozygosity in hybrid genomes and that the patterns observed in genomes from *C. albicans* strains are similar to those of hybrid strains of *C. orthopsilosis* clade 1 (e.g. strain MCO456), which underwent extensive levels of LOH after hybridization (Pryszcz et al. 2014; Schröder et al. 2016).

**Figure 1.**
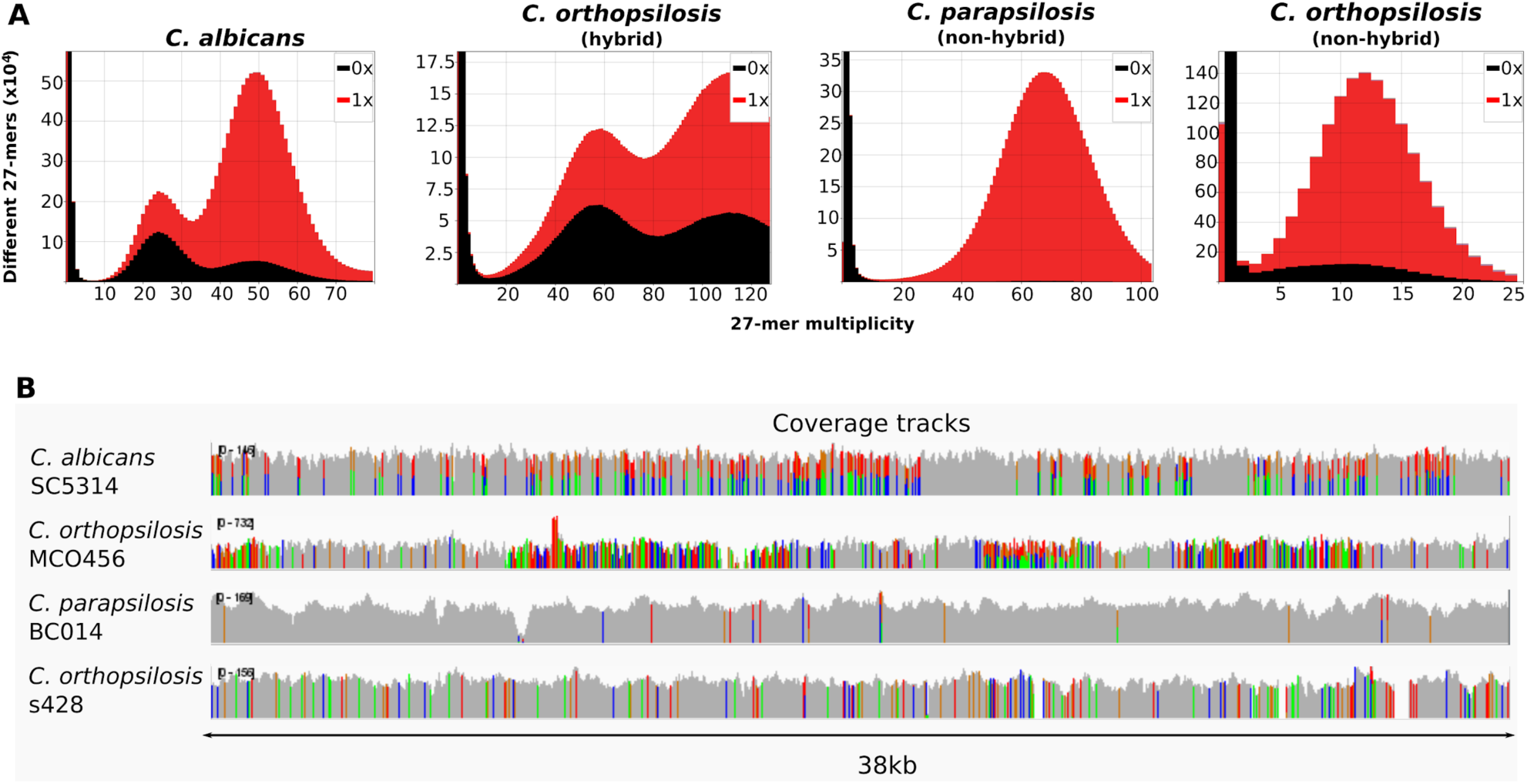
Comparison of the genomic patterns observed in *C. albicans* and hybrid and non-hybrid species. **A)** 27-mer frequency for SC5314 (*C. albicans*), MCO456 (*C. orthopsilosis* hybrid), BC014 (*C. parapsilosis* non-hybrid) and s428 (*C. orthopsilosis,* non-hybrid parental lineage A), and their respective presence (red) or absence (black) in the respective reference genome (plots were obtained with KAT (Mapleson et al. 2016)); **B)** Coverage tracks for illustrative genomic regions of the above mentioned genomic sequencing libraries, when aligned to the respective genomes. Colors indicate polymorphic positions. Positions with more than one color correspond to heterozygous variants. Visualizations were performed with IGV (Thorvaldsdóttir, Robinson, and Mesirov 2013).

### Heterozygosity patterns in *C. albicans* are comparable to those of hybrid lineages

We next assessed heterozygosity levels by aligning genomic reads of *C. albicans* strains, including putative wild strains isolated from oak trees (Bensasson et al. 2019), on the reference for haplotypes A and B, independently (see Materials and Methods). Similar results were obtained for the two haplotypes, and therefore we only report the results for haplotype A. This analysis revealed a genome-wide average heterozygosity of 6.70 heterozygous variants per kilobase (kb), which depending on the clade varied from 4.59 to 8.62 variants/kb (Table S1). Once again, these values are comparable to what was observed for *C. orthopsilosis* MCO456 (clade 1) strain, where we found 7.16 heterozygous variants/kb (Table S2). Of note, *C. albicans* strains isolated from oak trees were highly heterozygous, as previously reported (Bensasson et al. 2019), but their values were in the range of what we observed for clinical isolates.

Furthermore, as previously described (Hirakawa et al. 2015; Ropars et al. 2018; Wang, Bennett, and Anderson 2018; Bensasson et al. 2019), the heterozygous variants in *C. albicans* were not homogeneously distributed across the genome. Rather, these variants were concentrated in heterozygous regions separated by what appeared to be blocks of LOH (Figure 1B and Figure S2). Moreover, we could identify some regions which were highly heterozygous in some strains whereas in others corresponded to LOH regions that tended to alternative haplotypes (Figure S2). These patterns were reminiscent to the ones observed in *Candida* hybrid strains (Figure 1B and Figure S2) (Pryszcz et al. 2014; Pryszcz et al. 2015; Schröder et al. 2016; Mixão et al. 2019).

The comparison of heterozygosity patterns between different species is often hampered by the use of different methodologies and criteria to define heterozygous and homozygous regions in different studies. For instance, while studies performed so far on *C. albicans* distinguished heterozygous from LOH regions based on SNP density within windows of 5kb (Ene et al. 2018; Wang, Bennett, and Anderson 2018; Hirakawa et al. 2015), 10kb (Ropars et al. 2018), or even 100kb length (Bensasson et al. 2019), studies performed on hybrids defined these blocks based on the distance between heterozygous positions (Pryszcz et al. 2014; Pryszcz et al. 2015; Mixão et al. 2019). This last approach makes the boundaries between heterozygous and LOH blocks more flexible and precise and avoids averaging levels of heterozygosity when a window spans both homozygous and heterozygous regions. We therefore decided to use the methodological framework previously applied and validated in the study of hybrid species (see Materials and Methods) to analyze genome-wide heterozygosity patterns and define LOH blocks in *C. albicans* and compare them to patterns in established *Candida* hybrids.

On average, in *C. albicans* strains, we could define 7,059 heterozygous blocks and 16,492 LOH blocks, representing 11.18% and 85.74% of the genome, respectively. This is again notably similar to *C. orthopsilosis* hybrid clade 1, where 84.85% of the genome underwent LOH (Tables S2 and S3). Although these heterozygous blocks in *C. albicans* comprised most of the heterozygous SNPs (on average 59.35%), 12.79% of the heterozygous variants were placed within LOH blocks, and the remaining 27.86% in undefined regions (see Materials and Methods). The number of heterozygous variants outside heterozygous blocks was comparatively much higher than those found in *C. orthopsilosis* or *C. metapsilosis* hybrids (where it ranged from 2.38% to 5.81%, depending on the strain, Table S2), but notably closer to that found in the recently identified *C. inconspicua* hybrids (up to 23.23%, Mixão et al. 2019). The higher number of heterozygous variants within LOH blocks would suggest that *C. albicans* and *C. inconspicua* LOH blocks have been accumulating mutations for a longer time, as compared to *C. orthopsilosis* or *C. metapsilosis*. In addition, it is worth noting that the union of the heterozygous blocks of the 61 *C. albicans* strains analyzed in this work corresponded to more than half (53.17%, Table S4) of *C. albicans* genome, and this value is expected to increase with a larger sample size. Although from these analyses, we cannot completely exclude the possibility that the accumulation of mutations followed by recombination between different strains is responsible for the heterozygosity in *C. albicans* (Ropars et al. 2018; Wang, Bennett, and Anderson 2018), their strong similarity with what is observed for *C. inconspicua, C. orthopsilosis* and *C. metapsilosis* hybrid strains and the high level of heterozygosity of *C. albicans* genome strongly suggest a scenario where hybridization between two diverged lineages was followed by extensive LOH.

### The majority of heterozygous variants in *C. albicans* predate the diversification of known clades

The level of conservation of heterozygosity patterns across *C. albicans* strains of different clades can be used to assess the possibility of an ancestral hybridization event. Indeed, if a hybridization event between two divergent lineages, rather than the vertical accumulation of variants, was responsible for a sizable fraction of the heterozygosity levels observed across *C. albicans* genomes, then we would expect that a significant amount of heterozygous SNPs within heterozygous blocks would be shared by strains from deeply divergent clades, as their origin would have predated the diversification of the different *C. albicans* clades. To test this, we selected three non-overlapping sets of four strains (considered as replicates, see Materials and Methods) so that each set contains a representative strain of each of four deeply divergent clades (Figure 2A), according to the recent strain phylogeny described by (Ropars et al. 2018). For each group we compared the heterozygous positions in heterozygous and LOH regions shared by the different clades (Figure 2B, see Materials and Methods). We inferred that a heterozygous position in a given strain was ancestral if the most parsimonious reconstructed scenario (i.e. the one involving the lowest number of mutations) pointed to the same heterozygous genotype in the common ancestor of the four clades. The results, considering positions that could be unambiguously inferred, were consistent between the different groups (Figure 2C, Table S5), and showed that, on average, between 79.93% and 83.34% of the heterozygous positions within heterozygous blocks were ancestral (i.e. were present before the divergence of the clades), as compared to 15.28% to 20.10% of heterozygous positions within LOH blocks. Interestingly, the density of SNPs supporting a hybridization scenario presented a normal distribution with a peak at 20 SNPs/kb in all strains (Figure S3). The high level of shared SNPs in heterozygous regions across different clades was kept even when considering coding and non-coding regions separately (Table S5). These results strongly suggest that most of the heterozygous variants in heterozygous blocks were already present in a putative *C. albicans* ancestor, whereas most of the variants in LOH blocks appeared later, by independent accumulation in the different lineages.

**Figure 2.**
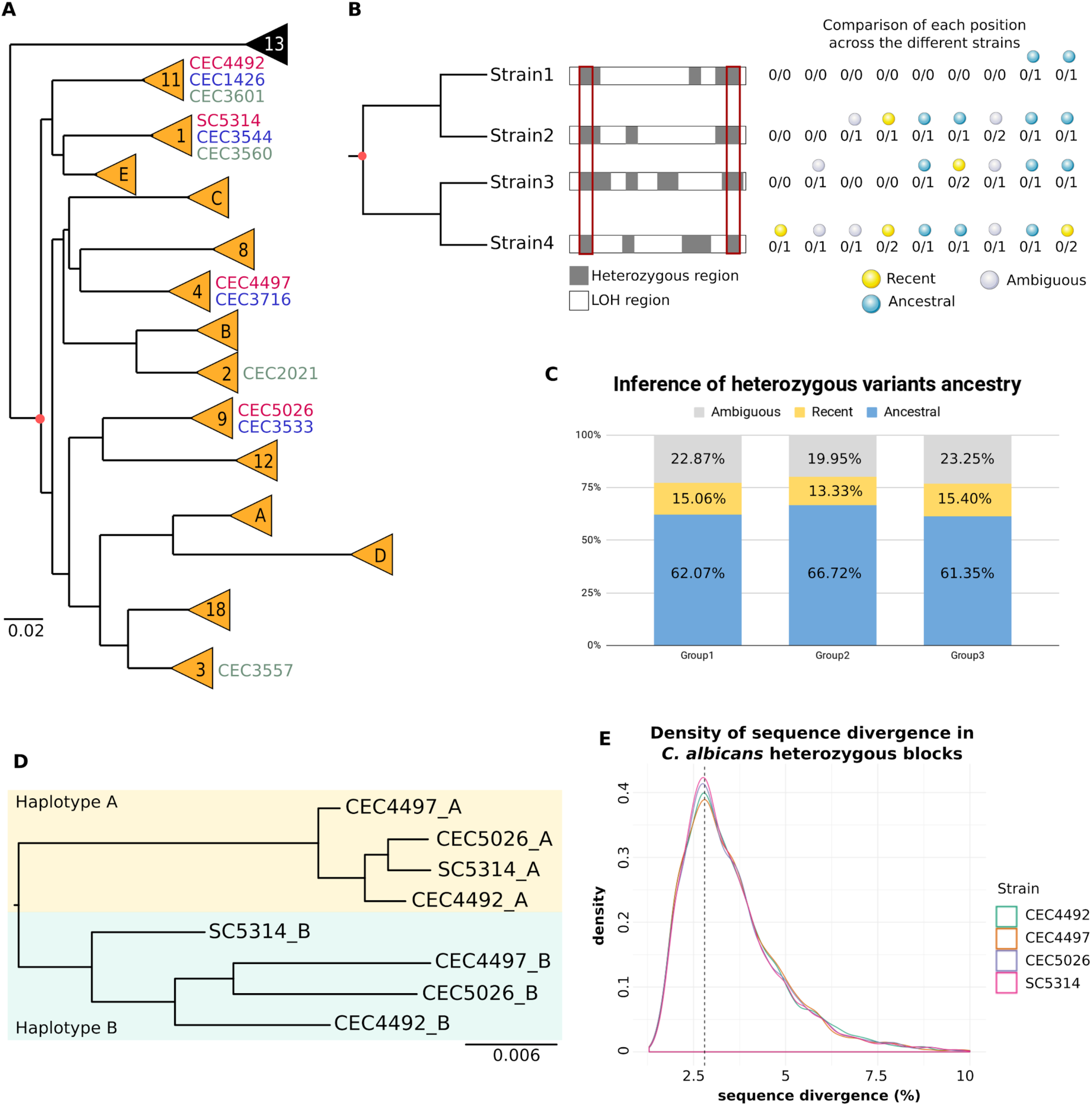
Analysis of *C. albicans* heterozygosity patterns. **A)** Schematic phylogenetic tree adapted from (Gabaldón and Fairhead 2019) indicating the different *C. albicans* clades in orange, and a potential common ancestor in red. Strains used for the comparison of heterozygous positions across different clades are indicated, with strains of the same group of analysis being written with the same color (red - group 1, blue - group 2 and grey - group 3); **B)** Schematic representation of the methodology for the comparison of heterozygous SNPs in heterozygous blocks (grey). The intersection of the heterozygous blocks is represented by the red rectangle (although not shown, the same approach was used for the analysis of LOH blocks). Examples of possible combinations of genotypes across the four strains are given (“0” - allele similar to the reference, “1” - allele different from the reference, “2” - allele different from the reference and from “1”). The most parsimonious path for the SNPs observed in each position was reconstructed. The decision taken for each position in a given strain is represented by yellow (recent), blue (ancestral) or grey (ambiguous) sferes; **C)** Plot of the average proportion ambiguously (grey) and unambiguously assigned positions for each group of strains. For unambiguously assigned positions, the proportion of recent and ancestral positions is shown in yellow and blue, respectively; **D)** Maximum likelihood phylogeny of the aligned reconstructed haplotypes A and B for the intersection of heterozygous blocks >100bp of group 1; **E)** Sequence divergence distribution in heterozygous blocks of *C. albicans* SC5314, CEC4492, CEC4497 and CEC5026.

In agreement with these results, a Maximum Likelihood phylogeny of the reconstructed haplotypes in heterozygous regions for the same groups of four strains (see Materials and Methods) indicated that the phylogenetic distance between the two haplotypes is higher than the distance between the different strains (Figure 2D and Figure S4). A similar approach was used in the past to confirm a hybrid origin of *C. orthopsilosis* (Pryszcz et al. 2014). Based on the number of variants per kilo-base in heterozygous blocks we estimated that the homeologous chromosomes are currently approximately 3.5% divergent at nucleotide level (Figure 2E and Table S3). This estimation was consistent across the sixty-one different strains (ranging from 3.44% to 3.59%, Table S3), and therefore unrelated to their overall level of heterozygosity. These analyses strongly suggest that most of the divergence between haplotypes in *C. albicans* heterozygous blocks was already present in a putative, highly heterozygous ancestor, with an estimated 2.8% divergence between the homeologous chromosomes (assuming ∼80% of the current variants in heterozygous blocks were heterozygous in the ancestor, in line with our estimations above). We consider that the most likely scenario to explain such pattern is a hybridization event between two divergent lineages.

### *Candida africana* is an evolved *C. albicans* lineage that underwent massive LOH, and not a putative parental of the hybrid ancestor of *C. albicans*

*C. africana* was proposed to be ranked as a species in 2001 (Tietz et al. 2001). However, although it presents particular phenotypes, the genetic similarity with *C. albicans* makes this controversial, and *C. africana* is often considered another *C. albicans* clade (clade 13, Romeo et al. 2013). In a recent population genomics study, Ropars et al. showed that, contrary to *C. albicans*, *C. africana* is highly homozygous, and hypothesized that massive LOH might have occurred in this clade (Ropars et al. 2018).

Given our results indicating that *C. albicans* originates from a hybridization event, we wanted to investigate the possibility that *C. africana* lineage could correspond to one of the parentals involved in the hybridization. To address this, we selected a sample of *C. africana* strains (see Materials and Methods) and analyzed the respective genomic patterns. As expected given the high levels of LOH previously described for this lineage (Ropars et al. 2018), *C. africana* presented low levels of heterozygosity, with an average of 2.68 heterozygous variants/kb, which were still high when compared to non-hybrid strains (Table S6). Therefore, we decided to define heterozygous regions in *C. africana* strains. In contrast to *C. albicans*, only 3.8% of the genomes, on average, corresponded to heterozygous blocks. These blocks have a current haplotype divergence of 3.7%, close to the 3.5% mentioned above for *C. albicans* (Table S6).

If *C. africana* was indeed one of *C. albicans* parents, we would expect the homozygous regions of a given chromosome to correspond always to the same *C. albicans* haplotype. The available phased genome of *C. albicans* is based on the heterozygous strain SC5314 (Muzzey et al. 2013), which already underwent LOH. Thus, only the regions of the phased reference genome corresponding to heterozygous regions of this strain can be used to assess distinct haplotypes in the inferred ancestral hybrid. From these regions we selected heterozygous positions that were considered ancestral in the above mentioned analyses and compared them to homozygous regions in *C. africana* strains (see Materials and Methods for details). Our results indicate that similar proportions of homozygous positions in *C. africana* could be mapped to each of the two haplotypes (55% A, and 45% B, Table S7). This strongly suggests that each *C. africana* chromosome is a mosaic of the two ancestral haplotypes, indicating it descents from the same hybrid ancestor as the other *C. albicans* clades. This shared ancestral hybridization scenario is reinforced by the fact that for the majority (99%) of the ancestral *C. albicans* heterozygous positions one of the alleles was represented in *C. africana*.

### Continuous gen flux from divergent lineages cannot explain consistent patterns found across strains

Taken together, our results show compelling evidence for a highly heterozygous genome in the ancestor of currently sampled *C. albicans* and *C. africana* clades. The presence of highly divergent haplotypes, and its consistency over the entire *C. albicans* genome can be best explained by an ancestral hybridization event between two distinct lineages and subsequent evolution through LOH. The alternative scenario of continuous introgression between divergent lineages could as well explain the presence of heterozygous regions in *C. albicans* strains, but could not explain the observed similarity of heterozygosis patterns across strains (Figure 3). Indeed, in a single hybridization scenario followed by divergence, extant heterozygous blocks in divergent strains are expected to present similar levels of sequence divergence, to share the same ancestral heterozygous SNPs, and to show some degree of overlap in their patterns of heterozygosity, as we have described. Moreover, we expect similar levels of heterozygosity and LOH between the strains from the same hybridization event (Schroder et al. 2016). Such consistent features of heterozygous blocks across divergent strains are difficult to explain by the accumulation of lineage-specific and independent events of introgression. Importantly, an ancestral hybridization scenario readily explains not only the heterozygosity patterns found in extant strains, but also could provide and explanation for the origin of other peculiar characteristics of *C. albicans* such as the absence of a standard sexual cycle, or its ubiquitous diploid nature, as discussed further below.

**Figure 3.**
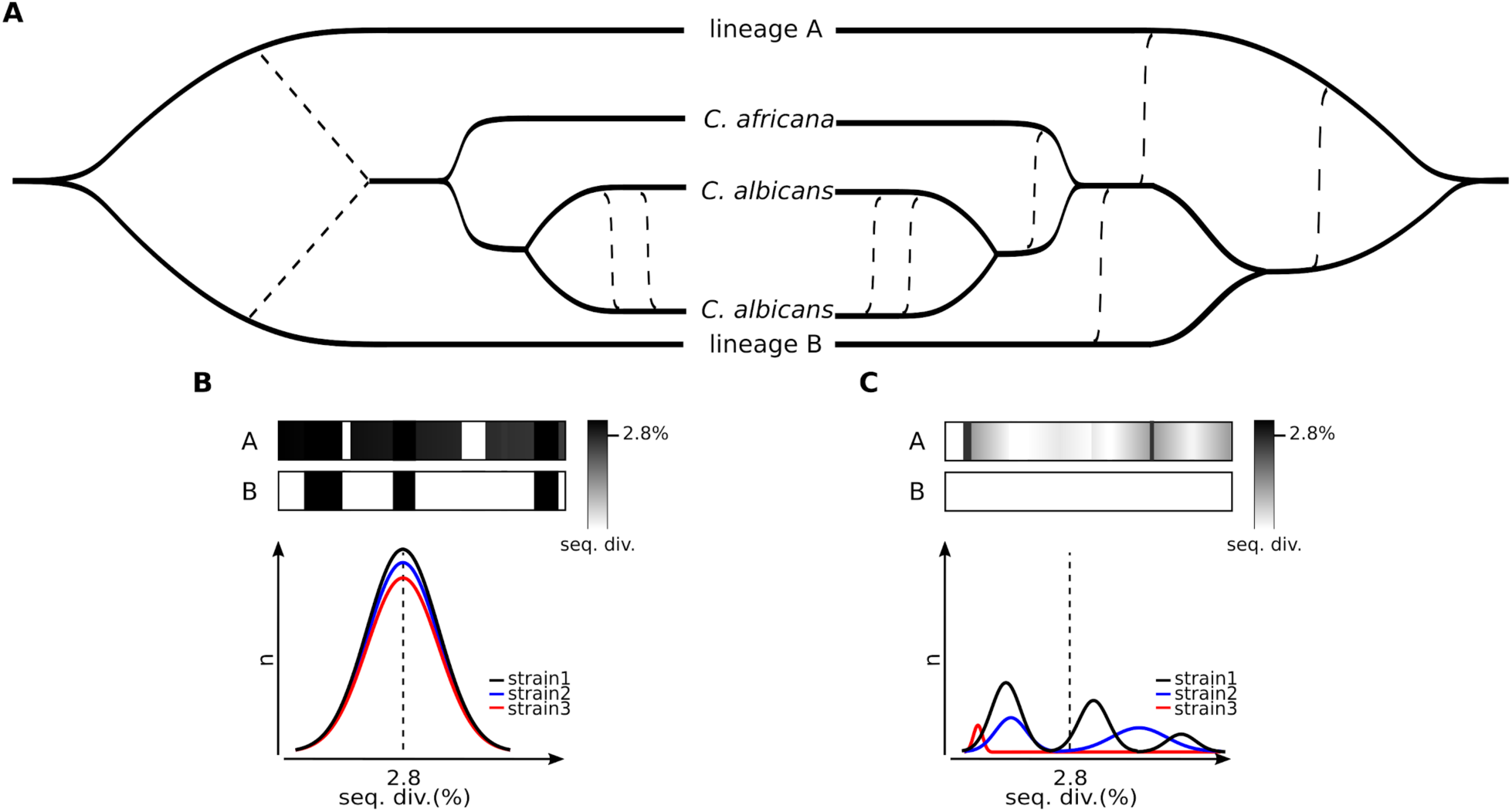
Schematic representation of plausible scenarios for the origin of *C. albicans* heterozygosity. **A)** The scenario proposed by this work is presented on the left, showing an ancestral hybridization event between two diverged lineages as the main source of *C. albicans* heterozygosity, followed by LOH, particularly extensive in *C. africana*, and more recent exchange of genomic material between *C. albicans* strains. The alternative scenario is presented on the right, showing inter-strain recombination as the only explanation for the heterozygous patterns observed in *C. albicans*. **B)** Scheme of the expected sequence divergence patterns if a single hybridization event was in the origin of the heterozygosity in *C. albicans*. After a hybridization event, the heterozygous blocks are expected to have similar sequence divergence, which is then reflected in a normal distribution. This divergence is expected to be similar in all strains originated from the same event. **C)** Scheme of the expected sequence divergence patterns if inter-strain recombination was the source of variability in *C. albicans*. In this scenario, the sequence divergence is time and strain dependent and therefore different patterns are expected between different blocks and between different strains.

## Discussion

Advances in Next-Generation sequencing have recently allowed the identification of many hybrid yeasts with clinical relevance (Pryszcz et al. 2014; Hagen et al. 2015; Pryszcz et al. 2015; Schröder et al. 2016; Mixão et al. 2019). Hybridization between diverged lineages is known to have an important role in the adaptation to new environments, or even in the emergence of new pathogens, as it is hypothesized to be the case of *C. metapsilosis* and *C. inconspicua* (Pryszcz et al. 2015; Mixão and Gabaldón 2018; Mixão et al. 2019). The hybrid nature of these strains was discovered by noting the presence of highly heterozygous genomes with a large divergence between alternative haplotypes and showing characteristic non-homogeneous distributions of heterozygous variants, resulting in highly heterozygous blocks separated by regions of low heterozygosity, likely resulting from LOH events. In most such cases, these hybrid strains are diploid and seem to be unable to undergo a normal sexual cycle (Pryszcz et al. 2014).

*C. albicans*, the most important yeast pathogen for human health (Barnett 2008), was previously shown to present genomic regions with high heterozygosity separated by what appeared to be blocks of LOH (Ropars et al. 2018). Para-sex and admixture were taken as the possible source for the observed levels of heterozygosity (Ropars et al. 2018; Wang, Bennett, and Anderson 2018). However, as the description of these patterns was reminiscent of that observed in hybrid lineages (Pryszcz et al. 2015; Schroder et al. 2016; Mixão et al. 2019), we hypothesized that hybridization could have been the initial source of genomic variability in *C. albicans*. Our results show compelling evidence that *C. albicans* descend from a hybrid between two divergent lineages. The genomic patterns observed in this pathogen are similar to those of other hybrids, specially *C. orthopsilosis* MCO456 and *C. inconspicua* (Pryszcz et al. 2014; Schröder et al. 2016; Mixão et al. 2019). This scenario clearly points to a highly heterozygous ancestor predating the divergence of currently known clades. The reconstructed most recent common ancestor of sequenced *C. albicans* strains would present most (53.17%) of its genome within heterozygous blocks. Current heterozygous regions present a 3.5% divergence between the two haplotypes, and we here infer that around 80% of such variants are ancestral, suggesting that the putative *C. albicans* ancestor had, at least, roughly 2.8% sequence divergence at the nucleotide level. From our analyses, we conclude that the existence of such level of divergence between two haplotypes in heterozygous regions can only be explained by these two haplotypes being genetically isolated for a long time. A hybridization between two previously isolated lineages, followed by LOH and further accumulation of SNPs would readily explain the observed patterns in *C. albicans*. Alternative scenarios accounting for the origin of heterozygous regions through independent introgression events would not explain the widespread presence of conserved heterozygous SNPs across strains from deeply divergent clades, nor the normal distribution observed for the levels of sequence divergence between haplotypes (Figure 2E).

We want to stress that the scenario of the hybridization between divergent lineages is agnostic to the consideration of the parental lineages as different or the same species. What is relevant is the realization of genomic chimerism predating the divergence of *C. albicans* clades. The species concept in microbes is controversial. The estimated 2.8% ancestral divergence between the hybridizing lineages largely exceeds levels of divergence found between most distantly related strains of well-studied yeast species such as *Sacharomyces cerevisiae*, where 1.1% sequence divergence was found between the most distantly related strains (Peter et al. 2018), and is higher than the estimated divergence between different described fungal species such as 1.4% between *Verticillium dahliae* and *Verticillium longisporum* D1 parental (Inderbitzin et al. 2011a; Inderbitzin et al. 2011b). On the other hand, it can be considered low when compared to ∼4.6% divergence between distant strains in *Saccharomyces paradoxus* (Liti, Barton and Louis 2006). Independently of the consideration of this putative ancestor as an inter- or intra-species hybrid, our results indicate that the ancestral hybrid did not backcrossed with any of the parental lineages but rather further evolved in a mostly clonal manner.

Inability to undergo meiosis and to complete a sexual cycle is a common feature of hybrids (Hunter et al. 1996; Wolfe 2015), including intra-species ones (Hou and Schacherer 2016; Rogers et al. 2018). Considering this, a hybridization scenario for the origin of *C. albicans* lineages would provide a plausible explanation for the origin of the inability of *C. albicans* to sporulate, or undergo a sexual cycle. In this particular case, we hypothesize that the improper chromosome pairing after hybridization and consequent impossibility of completing meiosis, contributed to the development of a parasexual cycle, an essential mechanism for *C. albicans* genomic plasticity. Alternatively, a pre-existing parasexual cycle may have facilitated the formation of hybrids by fusion of two unrelated diploid cells and subsequent concerted loss of chromosomes. In this regard, *C. tropicalis* have been shown to undergo a parasexual cycle under laboratory conditions (Seervai et al. 2013), although it is unclear how widespread is this ability among *Candida* species. This raises once more the question of the importance of hybridization for the emergence of yeast pathogens (Mixão and Gabaldón 2018) and poses the intriguing question of whether *C. albicans* ability to colonize and infect humans was an emerging phenotype enabled by this hybridization event.

## Materials and Methods

### NGS data selection

*C. albicans* paired-end reads used in this work are a subset of the data made publicly available under the BioProjects PRJNA432884 and PRJEB27862 (Ropars et al. 2018; Bensasson et al. 2019). Our sample was chosen based on different criteria. Specifically, we selected at least one strain from each SNP-based clade defined by Ropars et al., including *C. africana,* as this clade is highly homozygous (Ropars et. al 2018), and could correspond to a putative parental lineage. For clusters with more than one site of collection, one strain from each site was taken. In addition, the *C. albicans* type strain and two other environmental isolates were retrieved from (Bensasson et al. 2019). In the end, our sample consisted of a total of sixty-one *C. albicans* strains and eight *C. africana* (Table S1).

In order to compare our results with other species, we retrieved raw reads from Illumina paired-end sequencing libraries from known hybrid and non-hybrid strains from diverse *Candida* species. As representative of hybrid strains we selected *C. orthopsilosis* MCO456 (BioProject PRJEB4430, SRA ERX295059), s425, s433 and s498 strains (BioProject PRJNA322245, SRA SRX1776098, SRX1776103 and SRX1776124), and *C. metapsilosis* CP367 (BioProject PRJEB1698, SRA ERX221928) (Pryszcz et al. 2014; Pryszcz et al. 2015; Schröder et al. 2016). As representative of non-hybrid strains, we selected *C. orthopsilosis* s428 (BioProject PRJNA322245, SRA SRX1776102), three *C. parapsilosis* strains (BioProjects PRJEB1685 and PRJNA326748, SRA ERX221039, ERX221044 and SRX1875155) (Pryszcz et al. 2013; Schröder et al. 2016), and *C. tropicalis* ATCC200956 (BioProject PRJNA194439, SRA SRR868710).

### Library preparation and genome sequencing

As we considered important to compare *C. albicans* with the close related species *C. dubliniensis* and *C. tropicalis,* we decided to sequence two strains from our lab collections, namely 60-13 (*C. dubliniensis*) and CSPO (*C. tropicalis*). Genomic DNA extraction was performed using the MasterPure Yeast DNA Purification Kit (Epicentre, United States) following manufacturer’s instructions. Briefly, cultures were grown in an orbital shaker overnight (200 rpm, 30 °C) in 15 ml of YPD medium. Cells were harvested using 4.5 ml of each culture by centrifugation at maximum speed for 2 min, and then they were lysed at 65 °C for 15 min with 300 μl of yeast cell lysis solution (containing 1 μl of RNAse A). After being on ice for 5 min, 150 μl of MPC protein precipitation reagent were added into the samples, and they were centrifuged at 16.000 g for 10 min to pellet the cellular debris. The supernatant was transferred to a new tube, DNA was precipitated using 100% cold ethanol and centrifuging the samples at 16.000 g, 30 min, 4 °C. The pellet was washed twice with 70% cold ethanol and, once the pellet was dried, the sample was resuspended in 100 μl of TE. All gDNA samples were cleaned to remove the remaining RNA using the Genomic DNA Clean & Concentrator kit (Epicentre) according to manufacturer’s instructions. Total DNA integrity and quantity of the samples were assessed by means of agarose gel, NanoDrop 1000 Spectrophotometer (Thermo Fisher Scientific, United States) and Qubit dsDNA BR assay kit (Thermo Fisher Scientific).

Whole-genome sequencing was performed at the Genomics Unit from Centre for Genomic Regulation (CRG) with a HiSeq2500 machine. Libraries were prepared using the NEBNext Ultra DNA Library Prep kit for Illumina (New England BioLabs, United States) according to manufacturer’s instructions. All reagents subsequently mentioned are from the NEBNext Ultra DNA Library Prep kit for Illumina if not specified otherwise. 1 μg of gDNA was fragmented by ultrasonic acoustic energy in Covaris to a size of ∼600 bp. After shearing, the ends of the DNA fragments were blunted with the End Prep Enzyme Mix, and then NEBNext Adaptors for Illumina were ligated using the Blunt/TA Ligase Master Mix. The adaptor-ligated DNA was cleaned-up using the MinElute PCR Purification kit (Qiagen, Germany) and a further size selection step was performed using an agarose gel. Size-selected DNA was then purified using the QIAgen Gel Extraction Kit with MinElute columns (Qiagen) and library amplification was performed by PCR with the NEBNext Q5 Hot Start 2X PCR Master Mix and index primers (12–15 cycles). A further purification step were done using AMPure XP Beads (Agentcourt, United States). Final libraries were analyzed using Agilent DNA 1000 chip (Agilent) to estimate the quantity and check size distribution, and they were then quantified by qPCR using the KAPA Library Quantification Kit (KapaBiosystems, United States) prior to amplification with Illumina’s cBot. Libraries were loaded and sequenced 2 x 125 on Illumina’s HiSeq2500. Base calling was performed using Illumina pipeline software. In multiplexed libraries, we used 6-bp indexes. De-convolution was performed using the CASAVA software (Illumina, United States). Sequencing data are publicly available under the BioProject PRJNA555042.

### Raw sequencing data analysis

Raw sequencing data was inspected with FastQC v0.11.5 (http://www.bioinformatics.babraham.ac.uk/projects/fastqc/). Paired-end reads were filtered for quality bellow 10 or size bellow 31 bp and for the presence of adapters with Trimmomatic v0.36 (Bolger, Lohse, and Usadel 2014). The K-mer Analysis Toolkit (KAT, (Mapleson et al. 2016)) was used to get the 27-mers frequency and GC content of each library. This program was also used to inspect the representation of each 27-mer in the respective reference genome. The genome assemblies used as reference were: *C. albicans* assembly 22 (Muzzey et al. 2013), *C. orthopsilosis* 90-125 ASM31587v1 (Riccombeni et al. 2012), *C. metapsilosis* chimeric genome assembly (Pryszcz et al. 2015), and *C. parapsilosis* ASM18276v2 (Pryszcz et al. 2013).

### Read mapping and variant calling

Read mapping of each sequencing library to the respective reference genome assembly was performed with BWA-MEM v0.7.15 (Li, 2009). Picard integrated in GATK v4.0.2.1 (McKenna et al. 2010) was used to sort the resulting file by coordinate, as well as, to mark duplicates, create the index file, and obtain the mapping statistics. The mapping results were visually inspected with IGV version 2.4.14 (Thorvaldsdóttir, Robinson, and Mesirov 2013). Mapping coverage was determined with SAMtools v1.9 (Li et al. 2009).

Samtools v1.9 (Li et al. 2009) and Picard integrated in GATK v4.0.2.1 (McKenna et al. 2010) were used to index the reference and create its dictionary, respectively, for posterior variant calling. GATK v4.0.2.1 (McKenna et al. 2010) was used to call variants with the tool HaplotypeCaller set with -- genotyping-mode DISCOVERY --standard-min-confidence-threshold-for-calling 30 -ploidy 2. The tool VariantFiltration of the same program was used to filter the vcf files with the following parameters: -G-filter-name “heterozygous” -G-filter “isHet == 1” --filter-name “BadDepthofQualityFilter” -filter “DP <= 20 || QD < 2.0 || MQ < 40.0 || FS > 60.0 || MQRankSum < -12.5 || ReadPosRankSum < -8.0” --cluster-size 5 --cluster-window-size 20. In order to determine the number of SNPs/kb, a file containing only SNPs was generated with the SelectVariants tool. For this calculation only positions in the reference with 20 or more reads were considered for the genome size, and these were determined with bedtools genomecov v2.25.0 (Quinlan and Hall 2010).

### Heterozygous and homozygous blocks definition

To determine for each highly heterozygous strain the presence of heterozygous and LOH blocks, we adapted the procedure validated by Pryszcz et al. (Pryszcz et al. 2015). Briefly, bedtools merge v2.25.0 (Quinlan and Hall 2010) with a window of 100bp was used to define heterozygous regions, and by opposite, LOH blocks would be all non-heterozygous regions in the genome. The minimum LOH and heterozygous block size was established at 100bp. All regions that did not pass the requirements to be considered LOH or heterozygous blocks were classified as “undefined regions”. No filter for coverage was applied due to the low coverage of some libraries. For hybrid strains, the current divergence between the parentals was calculated dividing the number of heterozygous positions by the total size of heterozygous blocks.

### Comparison of SNPs across different strains

If *C. albicans* is a hybrid, we would expect that the majority of heterozygous SNPs in heterozygous blocks would be shared by the different strains, as this would mean that they were present before they diverged. On the other hand, in LOH blocks we would expect exactly the opposite, as the majority of heterozygous SNPs should correspond to new acquired mutations. To check this scenario, we compared the heterozygous and LOH blocks of four strains from different clades. Based on the phylogeny described by (Ropars et al. 2018), we decided to consider three groups of strains, which worked as replicates of the analysis (Figure 2A). Each group was comprised by two strains from each side of the first node of divergence of *C. albicans* strains. To ensure that the results were not influenced by events of recombination between different clades, we only selected strains that did not present signs of admixture with other clades according to Ropars et al. 2018. The first group of strains comprised CEC4492 (clade 11) and SC5314 (clade 1) from one side, and CEC4497 (clade 4) and CEC5026 (clade 9) from the other side. The second one comprised CEC1426 (clade 11) and CEC3544 (clade 1) from one side, and CEC3716 (clade 4) and CEC3533 (clade 9) from the other side. Finally, the third group comprised CEC3601 (clade 11) and CEC3560 (clade 1) from one side, and CEC2021 (clade 2) and CEC3557 (clade 3) from the other side. For each group, we obtained the intersection of their LOH blocks and the intersection of their heterozygous blocks using bedtools intersect v2.25.0 (Quinlan and Hall 2010). For each intersection, we inspected the heterozygous positions in each strain and compared them with the observed genotypes in the other three clades. For each position we reconstructed the most parsimonious scenario by assessing all possible mutational paths and choosing the one with the lower number of inferred mutations. When this scenario pointed to a similar heterozygous genotype in the common ancestor of the four strains, this position was assigned as “ancestral” for that strain. In case it would point to a different genotype, this heterozygous position was assigned as “recent”. When it was not possible to find a unique best scenario, the position was assigned as “ambiguous”. We also performed this analysis considering only blocks with an intersection >100bp. Furthermore, another analysis of both LOH and heterozygous blocks intersections was performed separately for coding and non-coding regions. For that, bedtools intersect v2.25.0 (Quinlan and Hall 2010) was used to obtain for each intersection the blocks with at least one overlap with coding regions annotated for *C. albicans* assembly 22 and available at Candida Genome Database (Skrzypek et al. 2017). Detailed information on the size of the intersection and positions considered for analyses are detailed in Table S5.

### Phylogenetic analysis considering the two haplotypes

The phylogenetic analysis of *C. albicans* considering the two haplotypes was performed individually for each of the mentioned groups of strains. Thus, for each group, we selected the intersection of heterozygous blocks (comprising the two haplotypes), which were defined as described above. Then, for each strain, the respective homozygous SNPs were substituted in the reference genome, and the two haplotypes separated according to their heterozygous SNP assignment (0/1), with the reference allele (0) being attributed to haplotype A, and the alternative (1) to haplotype B. Positions with INDELs in at least one of the strains were excluded. Positions with 1/2 SNPs were replaced by IUPAC ambiguity code. In the end, for each group we had an alignment of the two haplotypes of each strain. A Maximum-likelihood tree representative of each alignment was obtained with RAxML v8.2.8 (Stamatakis 2014), using the GTRCAT model. Midpoint rooting method was used to root the trees.

### Comparison of SNPs between *C. albicans* and *C. africana* strains

To assess whether *C. africana* was one of *C. albicans* parental lineages, we compared the heterozygous positions of *C. albicans*, with the homozygous positions of *C. africana.* As *C. albicans* genome was phased based on SC5314 (Muzzey et al. 2013), proved in this work to be a hybrid strain, only heterozygous regions in this strain are expected to represent the two parental haplotypes in the reconstructed phased genome. Therefore, for this analysis we considered the intersection of these positions with the heterozygous positions of each of the groups of *C. albicans* strains (check previous sections for details) and with the homozygous blocks defined for each *C. africana* strain. This analysis was performed independently for each group and each *C. africana* strain. Then, for each of these regions, we counted how many ancestral or recent positions identified in each of the four *C. albicans* strains of a given group (check previous sections for details) were shared (the same genotype was found in *C. albicans* and *C. africana*), or corresponded to a haplotype (haplotype A or B was present in both strains), or were undefined (none of the previous options was observed). Details on this analysis can be found in Table S7.

## Acknowledgments

This work was funded by the European Union’s Horizon 2020 research and innovation programme under the Marie Sklodowska-Curie grant agreement N° H2020-MSCA-ITN-2014-642095. TG group also acknowledges support from the Spanish Ministry of Economy, Industry, and Competitiveness (MEIC) for the EMBL partnership, and grants ‘Centro de Excelencia Severo Ochoa’ SEV-2012-0208, and BFU2015-67107 co-founded by European Regional Development Fund (ERDF); from the CERCA Programme/Generalitat de Catalunya; from the Catalan Research Agency (AGAUR) SGR857; and grants from the European Union’s Horizon 2020 research and innovation programme under the grant agreements ERC-2016-724173, and MSCA-747607. TG also receives support from a INB Grant (PT17/0009/0023 - ISCIII-SGEFI/ERDF). The authors thank Dr. Emilia Gomez, for providing CSPO strain, and all Gabaldón’s group, specially Susana Iraola, for laboratory work, and Marina Marcet-Houben, for the helpful discussions.

Sequencing data are publicly available under the BioProject PRJNA555042.

## Supplementary material

**Figure S1.** 27-mer frequency plots for SC5314, CEC4492, CEC4497, CEC5026 (*C. albicans*), MCO456, s425, s433, s498 (*C. orthopsilosis* hybrids), CP367 (*C. metapsilosis* hybrid), CBS6318, GA1, BC014 (*C. parapsilosis* non-hybrids), s428 (*C. orthopsilosis,* non-hybrid parental lineage A), 60-13 (*C. parapsilosis* non-hybrid), ATCC200956 and CSPO (*C. tropicalis* non-hybrid), and their respective presence (red) or absence (black) in the respective reference genome (plots were obtained with KAT (Mapleson et al. 2016)).

**Figure S2.** Coverage tracks for illustrative genomic regions of **A)** *C. albicans* strains; **B)** *C. albicans* strains with LOH towards different parents highlighted in the red box; **C)** *C. orthopsilosis* hybrid strains; and **D)** *C. metapsilosis* hybrid strain. Colors indicate polymorphic positions. Positions with more than one color correspond to heterozygous variants. Visualizations were performed with IGV (Thorvaldsdóttir, Robinson, and Mesirov 2013).

**Figure S3.** Distribution of ancestral SNPs/kb in group 1 (left), group 2 (center), and group 3 (right).

**Figure S4.** Maximum likelihood phylogeny of the aligned reconstructed haplotypes A and B for the intersection of heterozygous blocks >100bp of **A)** group 2; and **B)** group3.

**Table S1.** Detailed information on read coverage, mapping statistics and genomic variability of all *C. albicans* strains when aligned to the haplotype A.

**Table S2.** Detailed information on genomic variability of all analyzed non-*albicans* strains when aligned to the respective genome assemblies.

**Table S3.** Detailed information on the variability observed in heterozygous and LOH blocks for all *C. albicans* strains when mapped to haplotype A.

**Table S4.** Union and intersection of heterozygous blocks in each chromosome.

**Table S5.** Detailed results of the inference of heterozygous SNPs ancestrality for the three groups of *C. albicans* strains.

**Table S6.** Detailed information on read coverage, mapping statistics and genomic variability of all *C. africana* strains when aligned to the haplotype A of *C. albicans*.

**Table S7.** Detailed results of the inference of heterozygous SNPs ancestrality in comparison to *C. africana* for the three groups of *C. albicans* strains.

## References

Barnett JA. 2008. A History of Research on Yeasts 12: Medical Yeasts Part 1, *Candida albicans*. Yeast. 25(6):385–417.

Bennett RJ. 2015. The Parasexual Lifestyle of *Candida albicans*. Curr Opin Microbiol. 28(December):10–17.

Bensasson D, Dicks J, Ludwig JM, Bond CJ, Elliston A, Roberts IN, James SA. 2019. Diverse Lineages of Live on Old Oaks. Genetics. 211(1):277–88.

Berman J, Hadany L. 2012. Does Stress Induce (para)sex? Implications for *Candida albicans* Evolution. Trends Genet. 28(5):197–203.

Bolger AM, Lohse M, Usadel B. 2014. Trimmomatic: A Flexible Trimmer for Illumina Sequence Data. Bioinformatics. 30(15):2114–20.

Brown GD, Denning DW, Gow NAR, Levitz SM, Netea MG, White TC. 2012. Hidden Killers: Human Fungal Infections. Sci Transl Med. 4(165):165rv13.

Cauchie M, Desmet S, Lagrou K. 2017. *Candida* and Its Dual Lifestyle as a Commensal and a Pathogen. Res Microbiol. 168(9-10):802–10.

Ene IV, Farrer RA, Hirakawa MP, Agwamba K, Cuomo CA, Bennett RJ. 2018. Global Analysis of Mutations Driving Microevolution of a Heterozygous Diploid Fungal Pathogen. Proc Natl Acad Sci U S A. 115(37):E8688–97.

Forche A, Alby K, Schaefer D, Johnson AD, Berman J, Bennett RJ. 2008. The Parasexual Cycle in *Candida albicans* Provides an Alternative Pathway to Meiosis for the Formation of Recombinant Strains. PLoS Biol. 6(5):e110.

Gabaldón T, Carreté L. 2016. The Birth of a Deadly Yeast: Tracing the Evolutionary Emergence of Virulence Traits in *Candida glabrata*. FEMS Yeast Res. 16(2):fov110.

Gabaldón T, Fairhead C. 2019. Genomes Shed Light on the Secret Life of *Candida glabrata*: Not so Asexual, Not so Commensal. Curr Genet. 65(1):93–98.

Gladieux P, Ropars J, Badouin H, Branca A, Aguileta G, de Vienne DM, Rodríguez de la Vega RC, Branco S, Giraud T. 2014. Fungal Evolutionary Genomics Provides Insight into the Mechanisms of Adaptive Divergence in Eukaryotes. Mol Ecol. 23(4):753–73.

Hagen F, Khayhan K, Theelen B, Kolecka A, Polacheck I, Sionov E, Falk R, Parnmen S, Lumbsch HT, Boekhout T. 2015. Recognition of Seven Species in the *Cryptococcus gattii/Cryptococcus neoformans* Species Complex. Fungal Genet Biol. 78(May):16–48.

Hirakawa MP, Martinez DA, Sakthikumar S, Anderson MZ, Berlin A, Gujja S, Zeng Q, Zisson E, Wang JM, Greenberg JM, et al. 2015. Genetic and Phenotypic Intra-Species Variation in *Candida albicans*. Genome Res. 25(3):413–25.

Hou J, Schacherer J. 2016. Negative epistasis: a route to intraspecific reproductive isolation in yeast?. Curr Genet. 62(1):25–29.

Hunter N, Chambers SR, Louis EJ, Borts RH. 1996. The Mismatch Repair System Contributes to Meiotic Sterility in an Interspecific Yeast Hybrid. EMBO J. 15(7):1726–33.

Inderbitzin P, Bostock RM, Davis RM, Usami T, Platt HW, Subbarao KV. 2011a. Phylogenetics and Taxonomy of the Fungal Vascular Wilt Pathogen *Verticillium*, with the Descriptions of Five New Species. PloS One. 6(12):e28341.

Inderbitzin P, Davis RM, Bostock RM, Subbarao KV. 2011b. The Ascomycete *Verticillium longisporum* Is a Hybrid and a Plant Pathogen with an Expanded Host Range. PloS One. 6(3):e18260.

Jordà-Marcos R, Alvarez-Lerma F, Jurado M, Palomar M, Nolla-Salas J, León MA, León C, EPCAN Study Group. 2007. Risk Factors for Candidaemia in Critically Ill Patients: A Prospective Surveillance Study. Mycoses. 50(4):302–10.

Lass-Flörl C. 2009. The Changing Face of Epidemiology of Invasive Fungal Disease in Europe. Mycoses. 52(3):197–205.

Li H, unpublished data, https://arxiv.org/pdf/1303.3997.pdf, last accessed July 31, 2019

Li H, Handsaker B, Wysoker A, Fennell T, Ruan J, Homer N, Marth G, Abecasis G, Durbin R, 1000 Genome Project Data Processing Subgroup. 2009. The Sequence Alignment/Map Format and SAMtools. Bioinformatics. 25(16):2078–79.

Liti G, Barton DB, Louis EJ. 2006. Sequence Diversity, Reproductive Isolation and Species Concepts in Saccharomyces. Genetics. 174(2): 839–850.

Mapleson D, Garcia Accinelli G, Kettleborough G, Wright J, Clavijo BJ. 2016. KAT: A K-Mer Analysis Toolkit to Quality Control NGS Datasets and Genome Assemblies. Bioinformatics. 33(4):574–576.

McKenna A, Hanna M, Banks E, Sivachenko A, Cibulskis K, Kernytsky A, Garimella A, Altshuler D, Gabriel S, Daly M, DePristo MA. 2010. The Genome Analysis Toolkit: A MapReduce Framework for Analyzing next-Generation DNA Sequencing Data. Genome Res. 20(9):1297–1303.

Mixão V, Gabaldón T. 2018. Hybridization and Emergence of Virulence in Opportunistic Human Yeast Pathogens. Yeast. 35(1):5–20.

Mixão V, Perez Hansen A, Saus E, Boekhout T, Lass-Florl C, Gabaldón T. 2019. Whole-genome sequencing of the opportunistic yeast pathogen *Candida inconspicua* uncovers its hybrid origin. Front Genet. 10:383.

Muzzey D, Schwartz K, Weissman JS, Sherlock G. 2013. Assembly of a Phased Diploid *Candida albicans* Genome Facilitates Allele-Specific Measurements and Provides a Simple Model for Repeat and Indel Structure. Genome Biol. 14(9):R97.

OPATHY Consortium and Gabaldón T. 2019. Recent trends in molecular diagnostics of yeast infections: from PCR to NGS. FEMS Microbiol Rev. 43(5):517–547.

Peter J, De Chiara M, Friedrich A, Yue J-X, Pflieger D, Bergström A, Sigwalt A, Barre B, Freel K, Llored A, et al. 2018. Genome Evolution across 1,011 *Saccharomyces cerevisiae* Isolates. Nature. 556(7701):339–44.

Pfaller MA, Diekema DJ. 2007. Epidemiology of Invasive Candidiasis: A Persistent Public Health Problem. Clin Microbiol Rev. 20(1):133–63.

Pryszcz LP, Németh T, Gácser A, Gabaldón T. 2013. Unexpected Genomic Variability in Clinical and Environmental Strains of the Pathogenic Yeast *Candida parapsilosis*. Genome Biol Evol. 5(12):2382–92.

Pryszcz LP, Németh T, Gácser A, Gabaldón T. 2014. Genome Comparison of *Candida orthopsilosis* Clinical Strains Reveals the Existence of Hybrids between Two Distinct Subspecies. Genome Biol Evol. 6(5):1069–78.

Pryszcz LP, Németh T, Saus E, Ksiezopolska E, Hegedűsová E, Nosek J, Wolfe KH, Gacser A, Gabaldón T. 2015. The Genomic Aftermath of Hybridization in the Opportunistic Pathogen *Candida metapsilosis*. PLoS Genet. 11(10):e1005626.

Quinlan AR, Hall IM. 2010. BEDTools: A Flexible Suite of Utilities for Comparing Genomic Features. Bioinformatics. 26(6):841–42.

Riccombeni A, Vidanes G, Proux-Wéra E, Wolfe KH, Butler G. 2012. Sequence and Analysis of the Genome of the Pathogenic Yeast *Candida orthopsilosis*. PloS One. 7(4):e35750.

Rogers DW, McConnell E, Ono J, Greig D. 2018. Spore-autonomous fluorescent protein expression identifies meiotic chromosome mis-segregation as the principal cause of hybrid sterility in yeast. PloS Biol. 16(11):e2005066.

Romeo O, Tietz HJ, Criseo G. 2013. *Candida africana*: Is It a Fungal Pathogen?. Curr Fungal Infect Rep. 7:192.

Ropars J, Maufrais C, Diogo D, Marcet-Houben M, Perin A, Sertour N, Mosca K, Permal E, Laval G, Bouchier C, et al. 2018. Gene Flow Contributes to Diversification of the Major Fungal Pathogen *Candida albicans*. Nat Commun. 9(1):2253.

Schröder MS, Martinez de San Vicente K, Prandini THR, Hammel S, Higgins DG, Bagagli E, Wolfe KH, Butler G. 2016. Multiple Origins of the Pathogenic Yeast Candida orthopsilosis by Separate Hybridizations between Two Parental Species. PLoS Genet. 12(11):e1006404.

Seervai RNH, Jones Jr SK, Hirakawa MP, Porman AM, Bennett RJ. 2013. Parasexuality and Ploidy Change in *Candida tropicalis*. Eukaryot Cell. 12(12):1629–40.

Skrzypek MS, Binkley J, Binkley G, Miyasato SR, Simison M, Sherlock G. 2017. The Candida Genome Database (CGD): Incorporation of Assembly 22, Systematic Identifiers and Visualization of High Throughput Sequencing Data.” Nucleic Acids Res. 45(D1):D592–96.

Stamatakis A. 2014. RAxML Version 8: A Tool for Phylogenetic Analysis and Post-Analysis of Large Phylogenies. Bioinformatics. 30(9):1312–13.

Thorvaldsdóttir H, Robinson JT, Mesirov JP. 2013. Integrative Genomics Viewer (IGV): High-Performance Genomics Data Visualization and Exploration. Brief Bioinform. 14(2):178–92.

Tietz HJ, Hopp M, Schmalreck A, Sterry W, Czaika V. 2001. *Candida africana* sp. nov., a new human pathogen or a variant of *Candida albicans*?. Mycoses. 44(11-12):437–45.

Wang JM, Bennett RJ, Anderson MZ. 2018. The Genome of the Human Pathogen *Candida albicans* Is Shaped by Mutation and Cryptic Sexual Recombination. Mbio. 9(5):e01205–18.

Wolfe KH. 2015. Origin of the Yeast Whole-Genome Duplication. PLoS Biol. 13(8):e1002221.

