## Supplementary figures and images for "Genomic evidence for a hybrid origin of the yeast opportunistic pathogen *Candida albicans*"

### FigureS1.png

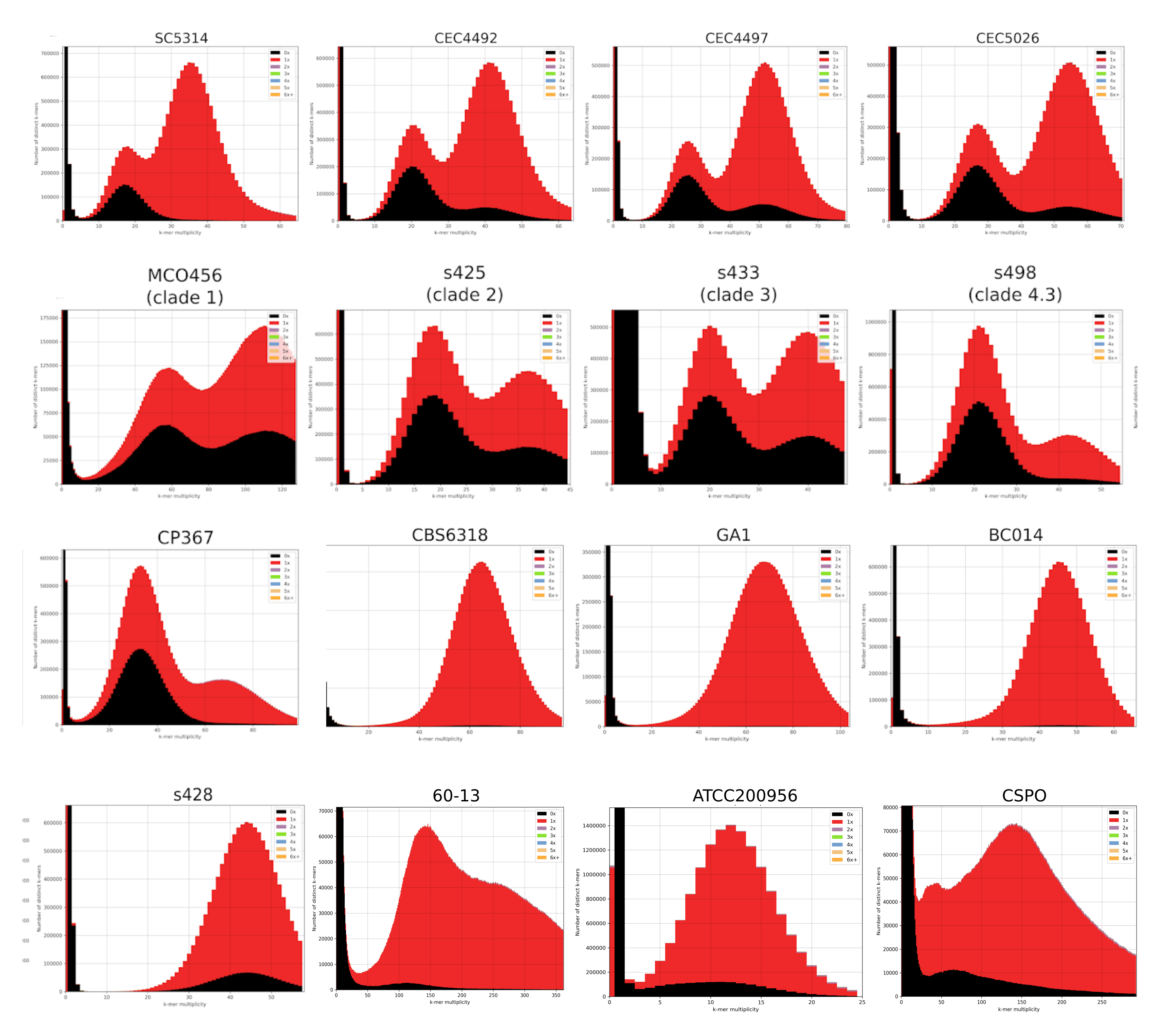

### FigureS2.png

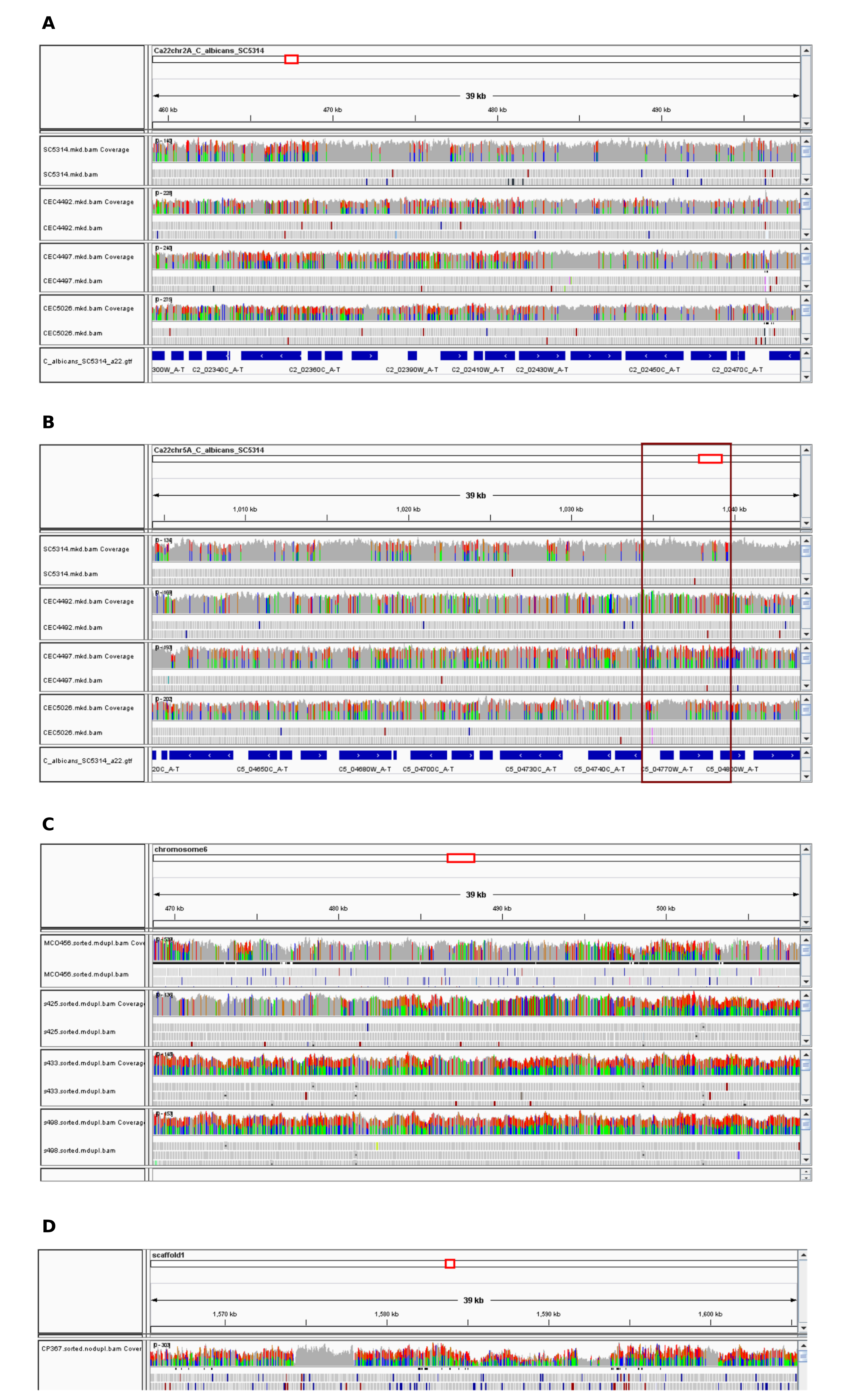

### FigureS3.png

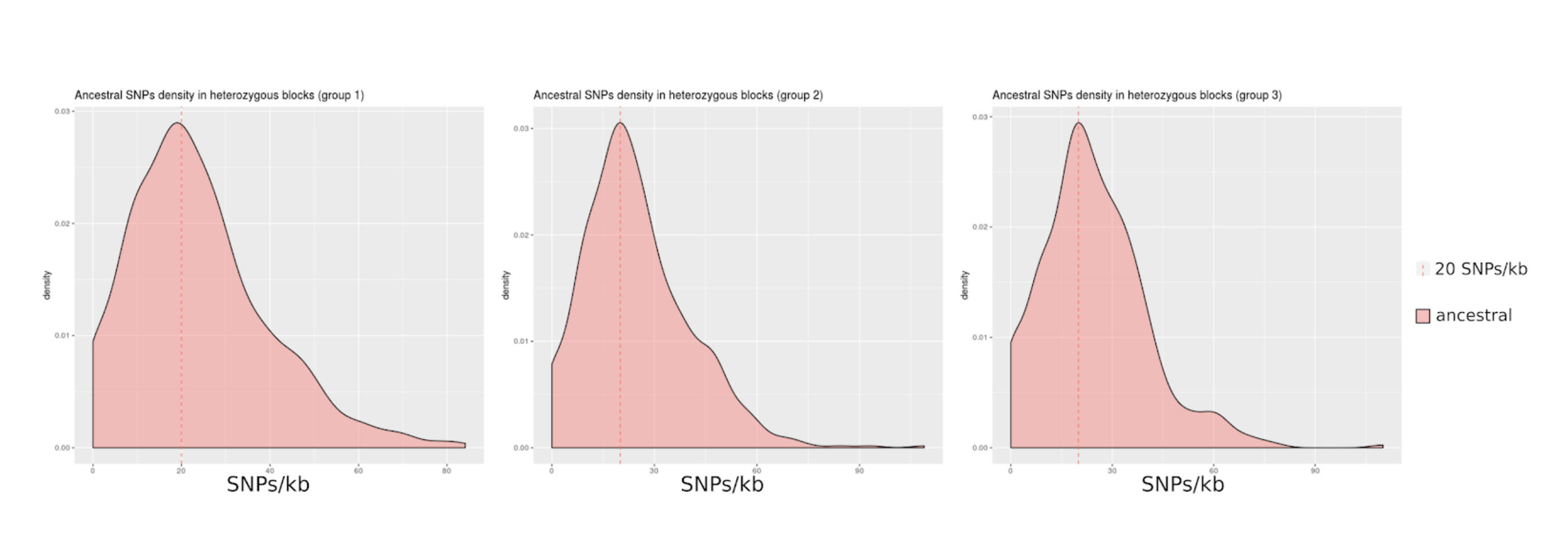

### FigureS4.png

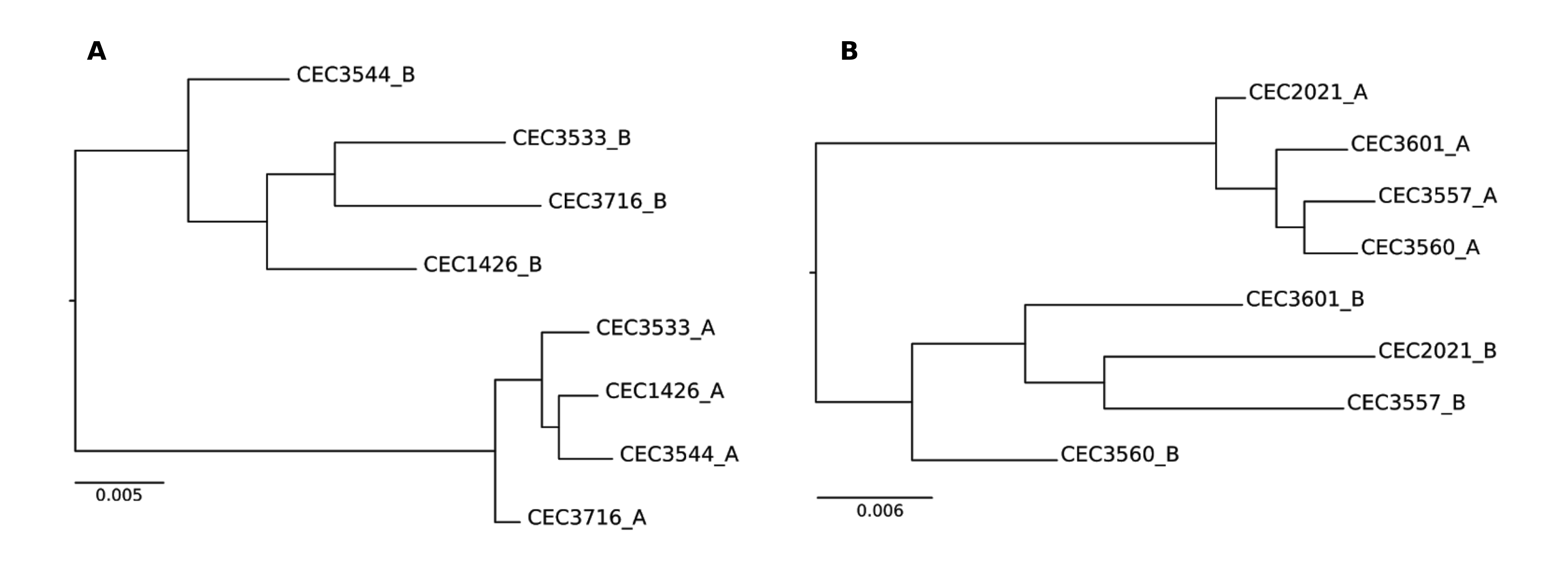
